# Longitudinal and stability-aware analysis reveals treatment-specific microRNA response signatures following immune-reconstitution and B-cell-targeted therapies in multiple sclerosis

**DOI:** 10.64898/2026.03.20.713174

**Authors:** Nasar Ata, Joshua S. Mytych, Mirela Cerghet, Ramandeep Rattan, Sumit Govil, Shailendra Giri, Yang Mao-Draayer

**Affiliations:** Department of Neurology, Henry Ford Health, Detroit, USA; School of Life and Basic Science Jaipur National University, Jaipur, India; Oklahoma Medical Research Foundation, Oklahoma City, OK, USA; Women’s Health Services, Henry Ford Health, Detroit, MI, 48202, USA

## Abstract

Disease-modifying therapies (DMT)s) for relapsing-remitting multiple sclerosis (RRMS) act through distinct immunological mechanisms, yet the within-patient molecular response programs associated with these therapies remain incompletely defined. Here, we reanalysed publicly available PBMC miRNA microarray data (GSE230064) using a longitudinal, robustness-focused framework to compare therapy-associated miRNA response patterns following cladribine versus ocrelizumab treatment. Baseline (t0) and 6-month post-treatment (t1) samples were paired within individuals and technical replicates consolidated prior to analysis, yielding a final paired cohort of 4 cladribine-treated and 6 ocrelizumab-treated patients. Within each treatment arm, we quantified per-patient Δ-miRNA (t1–t0) values and prioritized therapy-associated response features using a multi-evidence framework integrating effect direction, magnitude, directional consistency across individuals, and leave-one-out sensitivity. Cladribine treatment was associated with a highly coordinated, directionally concordant upregulation of five miRNAs including hsa-miR-27a-3p, hsa-miR-27b-3p, hsa-miR-503-5p, hsa-miR-148a-3p, and hsa-miR-26a-5p, all exhibiting 100% directional stability across patients and mean Δ-expression values ranging from +0.77 to +1.38. These miRNAs target pathways relevant to MS pathophysiology, including Th17/Treg balance, Wnt-β-catenin signaling, macrophage polarization, and epigenetic immune regulation. In contrast, ocrelizumab elicited a more selective response pattern, with five miRNAs including hsa-miR-100-5p, hsa-miR-410-3p, hsa-miR-432-5p, hsa-miR-296-5p, and hsa-miR-485-3p showing moderate directional stability (83%) and greater inter-individual heterogeneity, consistent with the more targeted mechanism of CD20+ B-cell depletion. Notably, the two treatment-associated signatures were non-overlapping, with hsa-miR-27b-3p representing the only miRNA shared with prior cross-sectional analyses of this dataset. The identified ocrelizumab-associated miRNAs implicate pathways including mTOR/IGF1R signaling, NF-κB regulation, RNA editing, and mitochondrial biogenesis, several of which are dysregulated in progressive MS. Together, these findings demonstrate that cladribine and ocrelizumab induce distinct, treatment-specific miRNA response architectures that reflect their divergent immunological mechanisms. This work establishes a stability-aware analytic template for extracting reproducible longitudinal miRNA signals from small paired RRMS cohorts and provides a ranked set of biologically plausible candidate miRNAs for prospective validation and mechanistic investigation.

## Introduction

Multiple sclerosis (Jakimovski et al., 2024) is a chronic, immune-mediated disorder of the central nervous system characterized by inflammation, neurodegeneration and demyelination(Arisi et al., 2023). Although numerous disease-modifying therapies (DMTs) are available, patients exhibit heterogeneous disease courses and treatment responses, so individualising therapy based on molecular stratification remains an unmet need(Arisi et al., 2023). MicroRNAs (miRNAs) are small (∼22 nucleotide) non-coding RNAs that regulate gene expression post-transcriptionally(Zabalza, Pappolla, Comabella, Montalban, & Malhotra, 2024). Their stability in blood, plasma and cerebrospinal fluid and ease of measurement make them attractive biomarkers for disease activity and therapeutic response. Deregulated miRNA expression patterns have been proposed as diagnostic tools in neuroinflammatory diseases such as MS.

Among high-efficacy DMTs used to treat MS, cladribine and ocrelizumab represent distinct mechanistic classes. Cladribine is a purine nucleoside analogue that must be phosphorylated to become active and is selectively taken up and activated by cells of lymphoid lineage. Because lymphocytes express high levels of the enzyme deoxycytidine kinase, cladribine accumulates in B and T cells, leading to DNA incorporation, induction of apoptosis and prolonged depletion of memory B cells while sparing much of the innate immune system. This pulsed therapy produces long-term immune reconstitution with a relatively favourable safety profile(Voo, Butzkueven, Stankovich, O’Brien, & Monif, 2021). In contrast, ocrelizumab is a humanized monoclonal antibody that binds the CD20 antigen on pre-B, mature and memory B cells (but not plasma cells or hematopoietic stem cells) and mediates depletion of these cells(Avasarala, 2017). Administration of ocrelizumab via intravenous infusion is approved for relapsing and primary-progressive MS; the therapy rapidly depletes circulating B cells and subsequently modulates other immune pathways, such as reducing inflammatory T-cell activity(Cencioni, Mattoscio, Magliozzi, Bar-Or, & Muraro, 2021).

Given the regulatory functions of miRNAs and their potential as biomarkers, characterising miRNA responses to cladribine and ocrelizumab could provide mechanistic insights and support personalised treatment strategies. However, most existing miRNA studies employ cross-sectional designs that cannot capture intra-individual dynamics. This study therefore utilises publicly available peripheral blood mononuclear cell miRNA data and a longitudinal, stability-aware analytical framework to identify treatment-specific miRNA signatures following cladribine-mediated immune reconstitution and ocrelizumab-mediated B-cell depletion

The initial study largely concentrated on cross-sectional comparisons of miRNA expression among treatment groups, whereas the current study re-evaluates the dataset utilizing a longitudinal within-subject approach. By calculating patient-specific Δ-miRNA profiles (t1−t0) and using directional consistency and leave-one-out stability measures, we uncover treatment-related miRNA signatures that are resilient to small cohort sizes and individual heterogeneity.

## Methods

### Dataset

The publicly accessible Gene Expression Omnibus (GEO) with accession number GSE230064 provided the miRNA expression dataset utilized in this investigation. PBMC samples from patients with relapsing-remitting multiple sclerosis (RRMS) receiving high-efficacy disease-modifying treatments, such as ocrelizumab (OCRE; n = 14) and cladribine (CLA; n = 11), as well as untreated control individuals (CTRL; n = 15), are included in the collection. While control participants were sampled just once, samples from treated patients were taken at baseline (t0) and six months after treatment started (t1). The original trial reported that ocrelizumab was given as 300 mg intravenous infusions twice over a two-week period, whereas cladribine pills were given at a cumulative dosage of 1.75 mg/kg over five days. The collection includes 65 microarray samples and 40 distinct participants overall, which reflects the inclusion of technical duplicates and repeated measurements throughout time. Only treated patients with full paired t0–t1 samples were kept for the current longitudinal study after technical replicates were collapsed, resulting in a final cohort of 10 paired patients (4 CLA, 6 OCRE).

### Study Cohort and Experimental Design

The dataset comprises peripheral blood mononuclear cell (PBMC) samples collected from individuals with relapsing-remitting MS (RRMS) treated with high-efficacy disease-modifying therapies, as well as untreated control subjects. RRMS patients received either Cladribine (CLA; *n* = 11 patients) or Ocrelizumab (OCRE; *n* = 14 patients). For treated patients, blood samples were collected at baseline prior to treatment initiation (t0) and again at 6 months post-treatment (t1). Untreated control subjects (*n* = 15) were sampled once at baseline. In total, 65 samples were included in the analysis. miRNA expression profiling was performed using an Agilent non-coding RNA microarray platform. Sample-level metadata, including treatment group, timepoint, tissue source, and cell type, were extracted directly from the GEO series matrix file. Although the GEO cohort included 11 cladribine-treated and 14 ocrelizumab-treated patients, only individuals with complete baseline (t0) and post-treatment (t1) samples were eligible for longitudinal Δ-miRNA analyses. After patient-level pairing and consolidation of technical replicates, the final paired cohort comprised 4 cladribine-treated and 6 ocrelizumab-treated patients. The statistical data were interpreted with caution due to the small number of matched patients that were included in each therapy arm within the study. Rather than strictly testing the hypothesis, the major objective of statistical testing was to prioritize the signals, and as a result, statistical significance was interpreted in conjunction with effect magnitude and stability criteria.

A detailed summary of sample characteristics and study design is provided in **Table 2**.

**Table 1.** Summary of dataset composition and sample selection from GEO dataset GSE230064.

| Category | Number |
| --- | --- |
| Total subjects | 40 |
| Cladribine-treated patients | 11 |
| Ocrelizumab-treated patients | 14 |
| Untreated control subjects | 15 |
| Total microarray samples (including technical replicates) | 65 |
| Baseline samples ($t_0$) | 33 |
| Follow-up samples ($t_1$) | 32 |
| Patients with complete paired samples used in longitudinal analysis | 10 |
| Cladribine paired patients | 4 |
| Ocrelizumab paired patients | 6 |

**Table 2.** Summary of study design and sample characteristics for cladribine- and ocrelizumab-treated patients included in the longitudinal Δ-miRNA analysis. The table reports sample counts, timepoints, and paired patient structure prior to consolidation of technical replicates.

| Characteristic | Cladribine | Ocrelizumab | Total |
| --- | --- | --- | --- |
| Total samples (including technical replicates of t0 and t1) | 29 | 36 | 65 |
| Baseline samples (t0) | 15 | 18 | 33 |
| Post-treatment samples (t1) | 14 | 18 | 32 |
| Patients with complete paired t0–t1 samples | 4 | 6 | 10 |
| Samples per patient (range) | 1–3 | 1–3 | — |
| Presence of technical replicates | Yes | Yes | Yes |
| Tissue source | Peripheral blood | Peripheral blood | — |
| Cell type | PBMC | PBMC | — |
| Expression platform | GEO microarray | GEO microarray | — |

### Quality Control and Preprocessing of miRNA Expression Data

Quality control (QC) analyses were performed to assess the global structure, distributional consistency, and technical integrity of miRNA expression data prior to downstream statistical analyses. miRNA expression values were extracted from the GEO series matrix file associated with accession GSE230064 and organized into a sample-by-feature expression matrix. No additional filtering, re-normalization, or batch correction was applied beyond the preprocessing performed by the original platform pipeline.

To make it easier to understand and keep the variance stable across the range of measured intensities, expression values were log2-transformed using log2(x + 1)(Lin, Du, Huber, & Kibbe, 2008). We used sample-wise density plots to look at the global expression distributions across samples. This helped us make sure that the expression profiles were consistent and find any technical outliers. We did unsupervised principal component analysis (PCA) on the log2-transformed expression matrix to find the main sources of variance and any possible sample-level artifacts or batch-driven structure. PCA was performed without the inclusion of phenotypic or treatment labels, guaranteeing an impartial evaluation of data quality(Blaise et al., 2021).

### Identification of Paired Samples and Handling of Technical Replicates

To enable longitudinal within-subject analyses, patient-level pairing of samples collected at baseline (t0) and 6 months post-treatment (t1) was performed for Cladribine- and Ocrelizumab-treated individuals. We got sample metadata from the GEO series matrix file. This included the treatment assignment, timepoint, and unique sample identifiers(Clough et al., 2024). A consistent patient-level identification was created by taking common prefixes from sample titles and using them to group together several technical measures that belonged to the same person. For every patient who received therapy, samples were organized by treatment arm and patient identifier. To be included in paired analyses, there had to be at least one baseline (t0) sample and one post-treatment (t1) sample. When there was more than one technical replicate for a patient and timepoint, the number of replicates was recorded to check for technical redundancy. All patients involved in subsequent analyses possessed complete t0–t1 pairings, and none were omitted due to absent timepoints.

### Technical Replicates and Computation of Longitudinal miRNA Changes

To measure changes in miRNA expression over time in the same patient after therapy, technical replicates that were taken at the same time and for the same patient were combined before the statistical analysis(Kahraman et al., 2018). We got the miRNA expression values from the processed expression matrix and changed them to log2(x + 1) to make the variance more stable and lower the effect of very high or low intensity values. For each patient i, miRNA j, and time point t (baseline t0 or post-treatment t1), we combined many technical replicates by finding their arithmetic mean(Kahraman et al., 2018). Formally, the collapsed expression value was defined as:

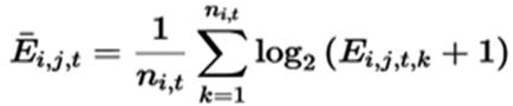

where Ei,j,t,kE_{i,j,t,k}Ei,j,t,k denotes the raw expression value of miRNA *j* for the *k*-th technical replicate of patient *i* at timepoint *t*, and ni,tn_{i,t}ni,t represents the number of available technical replicates for that patient and timepoint.

Longitudinal miRNA expression changes were then computed on a per-patient basis as the difference between post-treatment and baseline expression levels(Blaise et al., 2021)

where Δi,j\Delta_{i,j}Δi,j represents the within-subject change in expression of miRNA *j* for patient *i*. Positive Δ values indicate increased expression following treatment, whereas negative Δ values indicate decreased expression(Stevens, Herrick, Wolff, & Slattery, 2018). Δ-miRNA matrices were computed separately for Cladribine-treated and Ocrelizumab-treated patients and retained in comma-separated value (CSV) format for downstream statistical testing and visualization.

To identify treatment-associated miRNA changes, we performed statistical testing on within-subject Δ-miRNA values derived from paired baseline (t0) and post-treatment (t1) samples. For each patient and each miRNA, Δ-expression was computed as the difference between post-treatment and baseline expression levels (Δ = t1 − t0), following collapse of technical replicates by the mean on a log_2_(x+1) scale.

Separate analyses were performed for each treatment arm; Cladribine and Ocrelizumab, to prevent confounding molecular effects specific to each drug. A one-sample t-test was conducted for each miRNA within each treatment group to see if the mean Δ-expression substantially deviated from zero, in line with the null hypothesis of no systematic treatment-related change.

For each miRNA, the following statistics were computed: mean Δ-expression, standard deviation of Δ, t-statistic, and nominal p-value. Multiple-testing correction was performed using the Benjamini–Hochberg false discovery rate (FDR) procedure across all tested miRNAs within each treatment arm(Storey & Tibshirani, 2003). Due to the limited sample size and the presence of miRNAs with insufficient within-subject variance, FDR estimates were conservatively interpreted and reported primarily for transparency. All statistical analyses were implemented in Python using SciPy and stats models libraries.

### Stability and consistency analysis of treatment-associated Δ-miRNAs

To assess the robustness of treatment-associated Δ-miRNAs, a stability and consistency analysis was performed using subject-level Δ-miRNA matrices. Analyses were conducted separately for cladribine and ocrelizumab treatment arms.

We measured directional consistency for each miRNA by finding the percentage of patients whose Δ-expression was in the same direction as the group-level mean Δ. This statistic gives an easy-to-understand way to tell if changes seen in therapy are common to all participants or only to a few(Navon et al., 2009). We also did a leave-one-out (LOO) sensitivity analysis by taking away one individual at a time and calculating the mean Δ-expression again(Stevens et al., 2018). We looked just how sensitive group-level estimations are to individual participants by looking at how different LOO means were. Before using biological interpretation, these extra measures were utilized to check the stability of the top-ranked Δ-miRNAs.

We used a multi-evidence framework to combine the data to come up with strong treatment-related miRNA signatures. For each treatment arm (cladribine and ocrelizumab), miRNAs were maintained only if they exhibited: Consistent longitudinal change (Δ = t1 − t0), (ii) high ranking based on impact size, and (iii) substantial directional stability across individuals. This integrated method puts more value on consistency and repeatability than on standalone statistical significance. It produces treatment-specific miRNA signatures that show sustained longitudinal molecular responses.

## Results

Several additional quality control (QC) investigations were carried out to assess the overall quality and global structure of the miRNA expression data. All samples showed substantially overlapping distributions in sample-wise density plots of log2-transformed expression levels (Ben-Elazar et al., 2021)suggesting that there was no significant technical bias and that global expression patterns were consistent (**Figure 1A**). The dataset’s general comparability was supported by the fact that no sample showed an aberrant density pattern or a significantly altered distribution.

**Figure 1:**
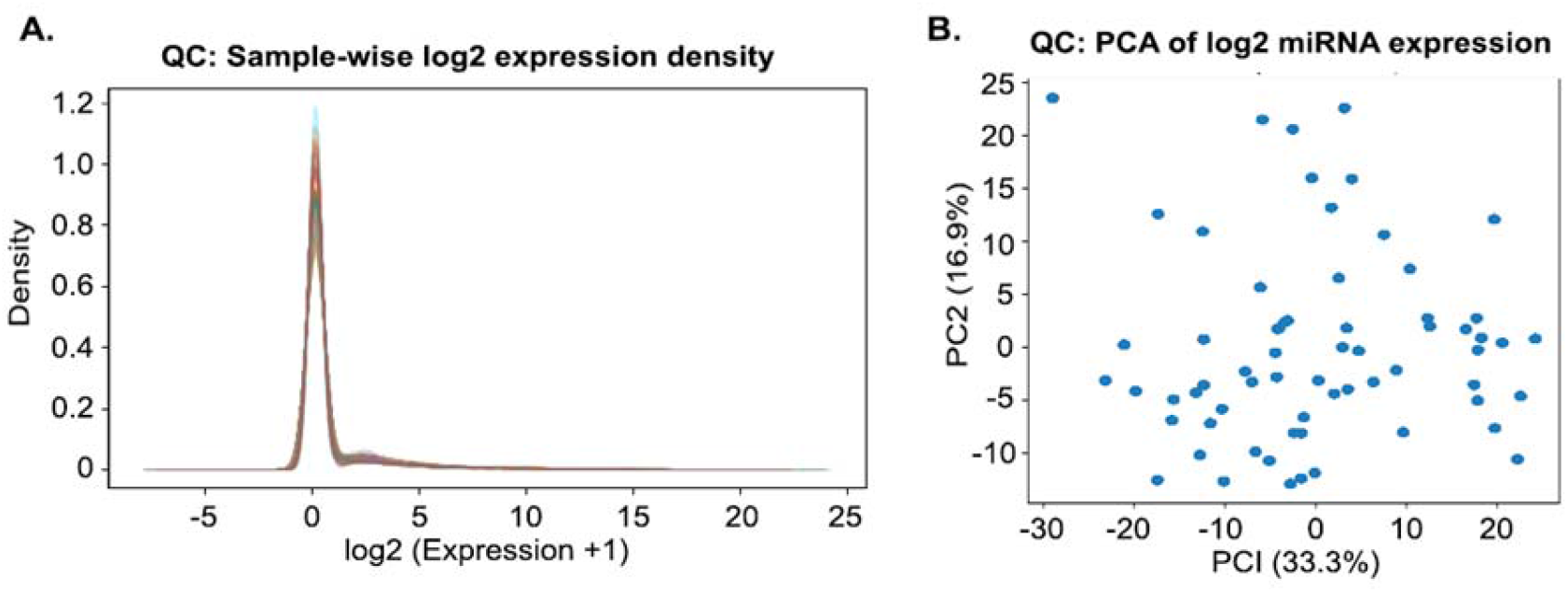
**(A)** Sample-wise density distributions of log2-transformed miRNA expression values across all samples, showing highly comparable global expression profiles. **(B)** Unsupervised principal component analysis (PCA) of log2-transformed miRNA expression values, demonstrating absence of extreme outliers or dominant technical effects.

The log2-transformed expression matrix was subjected to unsupervised principal component analysis (PCA), as previously demonstrated(Yang, Liu, Lu, Riggs, & Wu, 2017), which showed a continuous distribution of samples along the first two principal components. PC1 and PC2 accounted for a substantial proportion of the total variance, and was without evidence of dominant technical effects (Leek et al., 2010) **(Figure 1B)**. Crucially, neither a dominating technical axis nor any individual samples showed up as severe outliers as previously seen(Chen, Zhang, Wang, Bonni, & Zhao, 2020), indicating that batch effects or technical abnormalities were not the primary causes of global variation. The consistency of median expression levels and interquartile ranges across samples was further supported by the overlapping distributional characteristics observed across samples.

### Patient-Level Pairing and Longitudinal Sample Structure

Patient-level pairing identified a subset of treated individuals with complete longitudinal sampling suitable for within-subject analysis. From the full treated cohort, only patients with complete paired baseline and post-treatment samples were retained for longitudinal Δ-miRNA analyses, resulting in 4 cladribine-treated and 6 ocrelizumab-treated individuals. Six patients in the group treated with ocrelizumab satisfied the same requirements, whereas four patients in the group treated with cladribine had full baseline and post-treatment sample pairs. At least one technical replication was present for each timepoint in all paired patients, and few patients had more than one technical measurement at baseline and follow-up.

The number of patients with complete pairs and the existence of technical replicates at each timepoint are highlighted in **Table 3**, which provides an overview of the paired sample structure.

**Table 3.**
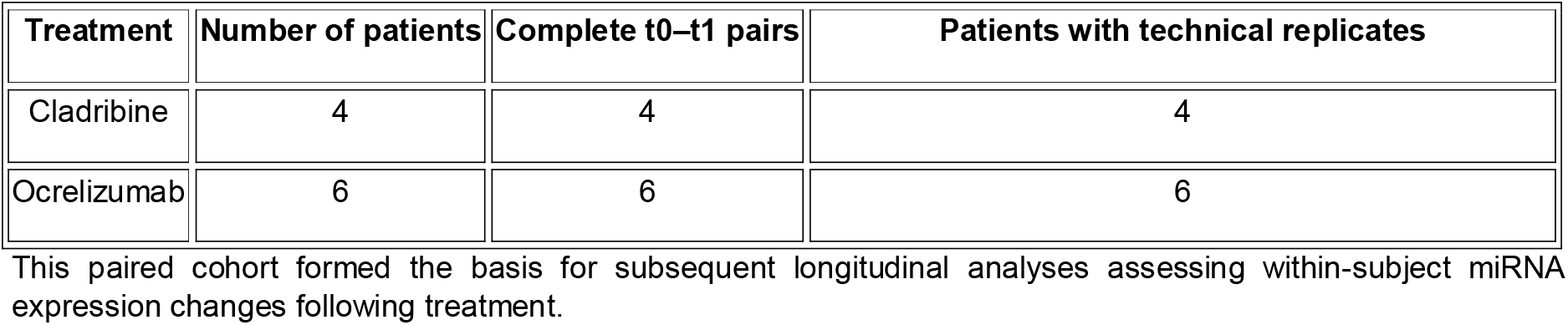
Summary of paired samples included in longitudinal analyses.

### Δ-miRNA profiles following cladribine and ocrelizumab treatment

Volcano plot analysis of Δ-miRNA values following cladribine treatment revealed a pronounced and structured pattern of miRNA changes **(Figure 2A)**. The volcano plot for cladribine (Figure 2A) shows strong changes in many miRNAs after treatment. Several miRNAs, including *hsa-miR-503-5p, hsa-miR-26a-5p, hsa-miR-27a-3p, hsa-miR-148a-3p*, and *hsa-miR-27b-3p*, show high increases in expression with strong significance. Most of the important changes are in the positive direction, while fewer miRNAs show decreases. Overall, this indicates that cladribine causes a broad and strong miRNA response. Large positive Δ-expression values with significant statistical support were shown by a group of miRNAs(Arisi et al., 2023), which clearly separated from the background distribution that was centred around zero. The overall distribution indicated a coherent treatment-associated shift rather than random variability. The volcano plot for ocrelizumab **(Figure 2B**) shows that most miRNAs do not change strongly after OCR treatment, as they are clustered near the centre with low significance. However, a few miRNAs, such as *hsa-miR-100-5p, hsa-miR-410-3p*, and *hsa-miR-432-5p*, show moderate increases in expression along with relatively higher significance. There are very few miRNAs showing strong decreases, indicating that the changes are mainly in one direction (increase). Overall, this suggests that ocrelizumab leads to selective and mild changes in specific miRNAs, rather than a broad or strong response across many miRNAs.

**Figure 2:**
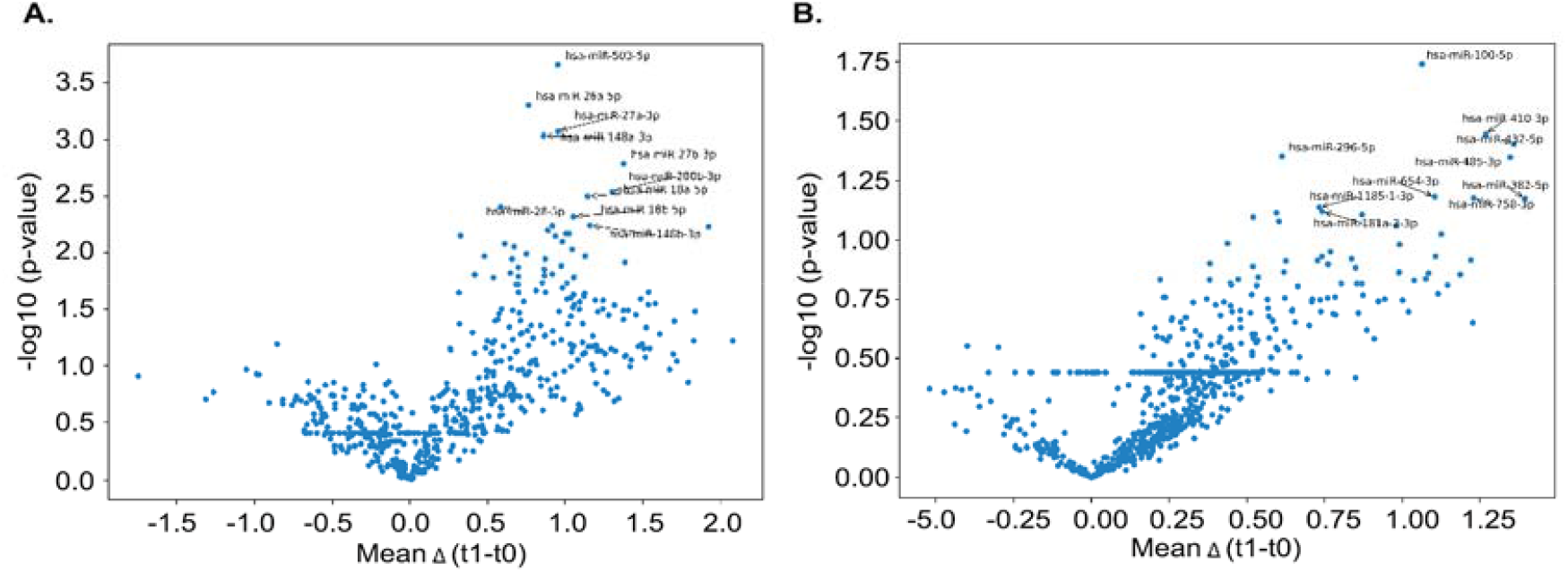
**(A)** Δ-miRNA volcano plot following cladribine treatment. **(B)** Δ-miRNA volcano plot following ocrelizumab (OCRE) treatment. Volcano plots show within-subject miRNA expression changes following cladribine or ocrelizumab treatment. All data points correspond to individual miRNAs, displayed by mean Δ-expression (t1 − t0) versus −log10(p-value). Representative miRNAs exhibiting the strongest treatment-associated changes are labelled. Fewer miRNAs exhibited large effect sizes, and the overall spread of Δ-expression values was narrower following cladribine treatment. Nevertheless, the volcano plot demonstrated a directional trend, indicating selective miRNA modulation following therapy. In contrast, the Δ-miRNA profile following ocrelizumab treatment showed a more modest pattern of change.

### Top treatment-associated Δ-miRNAs

A concise summary of the representative miRNAs showing the strongest longitudinal changes and statistical support for each treatment group is provided in **Table 4**. This table presents representative miRNAs with the most significant Δ-expression values and the strongest statistical validation within each group offering a concise summary of treatment-related signals. Together with the volcano plots, these findings demonstrate distinct molecular response patterns between cladribine and ocrelizumab at the miRNA level, similar to conclusions from the initial paper.

**Table 4.** Selected Δ-miRNAs associated with cladribine and ocrelizumab treatment: summarizing representative miRNAs showing treatment-associated changes following cladribine and ocrelizumab (OCRE) therapy. Mean Δ-expression (t1 − t0), standard deviation, test statistic, and nominal p-values are reported for each miRNA; false discovery rate (FDR) values are provided for transparency but interpreted conservatively due to limited sample size.

| Treatment | miRNA | Mean $\Delta$ ($t_1-t_0$) | SD $\Delta$ | t-statistic | p-value | FDR (q-value) |
| --- | --- | --- | --- | --- | --- | --- |
| CLA | hsa-miR-503-5p | 0.952332 | 0.088553 | 21.508778 | $2.20 \times 10^{-4}$ | 0.134808 |
| CLA | hsa-miR-26a-5p | 0.765690 | 0.093851 | 16.317194 | $5.01 \times 10^{-4}$ | 0.141283 |
| CLA | hsa-miR-27a-3p | 0.955557 | 0.139707 | 13.679444 | $8.45 \times 10^{-4}$ | 0.141283 |
| CLA | hsa-miR-148a-3p | 0.861446 | 0.129696 | 13.284068 | $9.22 \times 10^{-4}$ | 0.141283 |
| CLA | hsa-miR-27b-3p | 1.376618 | 0.250515 | 10.990301 | $1.61 \times 10^{-3}$ | 0.197760 |
| OCRE | hsa-miR-100-5p | 1.063450 | 0.754590 | 3.452085 | $1.82 \times 10^{-2}$ | 0.752265 |
| OCRE | hsa-miR-410-3p | 1.267997 | 1.090543 | 2.848071 | $3.59 \times 10^{-2}$ | 0.752265 |
| OCRE | hsa-miR-432-5p | 1.359277 | 1.201364 | 2.771462 | $3.93 \times 10^{-2}$ | 0.752265 |
| OCRE | hsa-miR-296-5p | 0.612689 | 0.562015 | 2.670351 | $4.43 \times 10^{-2}$ | 0.752265 |
| OCRE | hsa-miR-485-3p | 1.347491 | 1.239347 | 2.663228 | $4.47 \times 10^{-2}$ | 0.752265 |

**Table 5:** Final robust treatment-associated Δ-miRNA signatures derived from integrated longitudinal analysis summarizing miRNAs retained as robust treatment-associated signatures following integration of longitudinal change (Δ = t1 − t0), effect-size ranking, and directional stability across patients. Mean Δ-expression and directional stability percentages are shown separately for cladribine and ocrelizumab (OCRE) treatment groups.

| Treatment | miRNA | Mean $\Delta$ (t1–t0) | Directional Stability |
| --- | --- | --- | --- |
| CLA | hsa-miR-503-5p | +0.95 | 100 |
| CLA | hsa-miR-26a-5p | +0.77 | 100 |
| CLA | hsa-miR-27a-3p | +0.96 | 100 |
| CLA | hsa-miR-148a-3p | +0.86 | 100 |
| CLA | hsa-miR-27b-3p | +1.38 | 100 |
| OCRE | hsa-miR-100-5p | +1.06 | 83 |
| OCRE | hsa-miR-410-3p | +1.27 | 83 |
| OCRE | hsa-miR-432-5p | +1.36 | 83 |
| OCRE | hsa-miR-296-5p | +0.61 | 83 |
| OCRE | hsa-miR-485-3p | +1.35 | 83 |

### Ranking of Δ-miRNAs following cladribine and ocrelizumab treatment

Ranking of Δ-miRNAs following cladribine treatment, as previously done, we identified a subset of miRNAs exhibiting the largest and most consistent expression changes across patients (Arisi et al., 2023) (**Figure 3A**). These miRNAs showed high mean Δ-expression values, reinforcing the structured response pattern observed in the volcano analysis(Fissolo et al., 2023). Several of the top-ranked miRNAs belonged to related regulatory families, suggesting coordinated transcriptional modulation rather than isolated effects. A number of the cladribine-associated miRNAs are members of similar regulatory families, such as immune-regulatory miRNA clusters and the miR-27 family, indicating coordinated control of immune signalling pathways. The biological plausibility of the noted longitudinal alterations is strengthened by such family-level trends.

**Figure 3:**
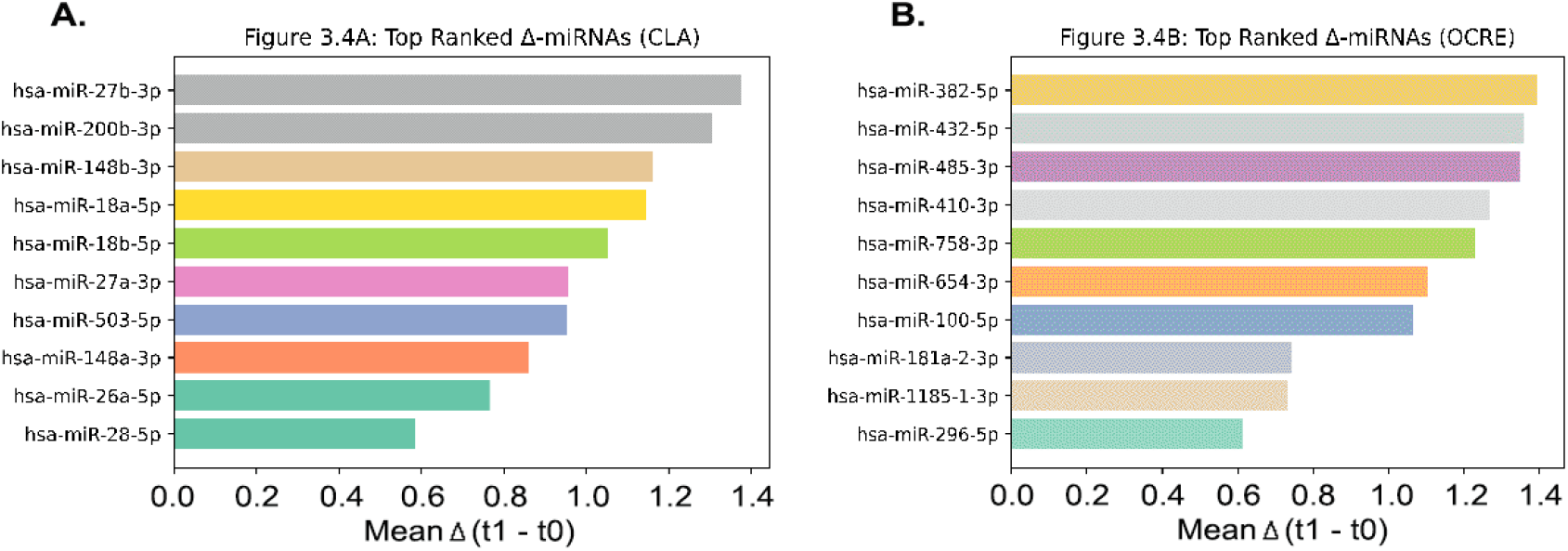
**(A)** Top-ranked Δ-miRNAs following cladribine treatment, ordered by mean within-subject Δ-expression (t1 − t0) across paired patients. **(B)**Top-ranked Δ-miRNAs following ocrelizumab (OCRE) treatment, ordered by mean Δ-expression (t1 − t0). Bars represent the magnitude of mean within-subject change for each miRNA.

In contrast, ranking of Δ-miRNAs following ocrelizumab(B. De Felice et al., 2025; Torres-Iglesias et al., 2025) treatment revealed a more selective response profile **(Figure 3B)**. The top-ranked miRNAs demonstrated moderate but consistent Δ-expression changes, reflecting a subtler molecular response compared with cladribine(Arisi et al., 2023). Despite the reduced magnitude, the ranked miRNAs highlight specific treatment-associated signals that are also unique to the specific treatment, no top miRNAs were found to overlap.

### Stability of Δ-miRNAs following cladribine and ocrelizumab treatment

Stability analysis revealed a high degree of directional consistency among the top-ranked Δ-miRNAs following cladribine treatment **(Figure 4A)**. Most miRNAs demonstrated consistent directionality across all or nearly all patients, indicating that observed Δ-expression changes were not driven by single individuals. This pattern indicates high directional consistency among cladribine-associated Δ-miRNAs across patients.

**Figure 4:**
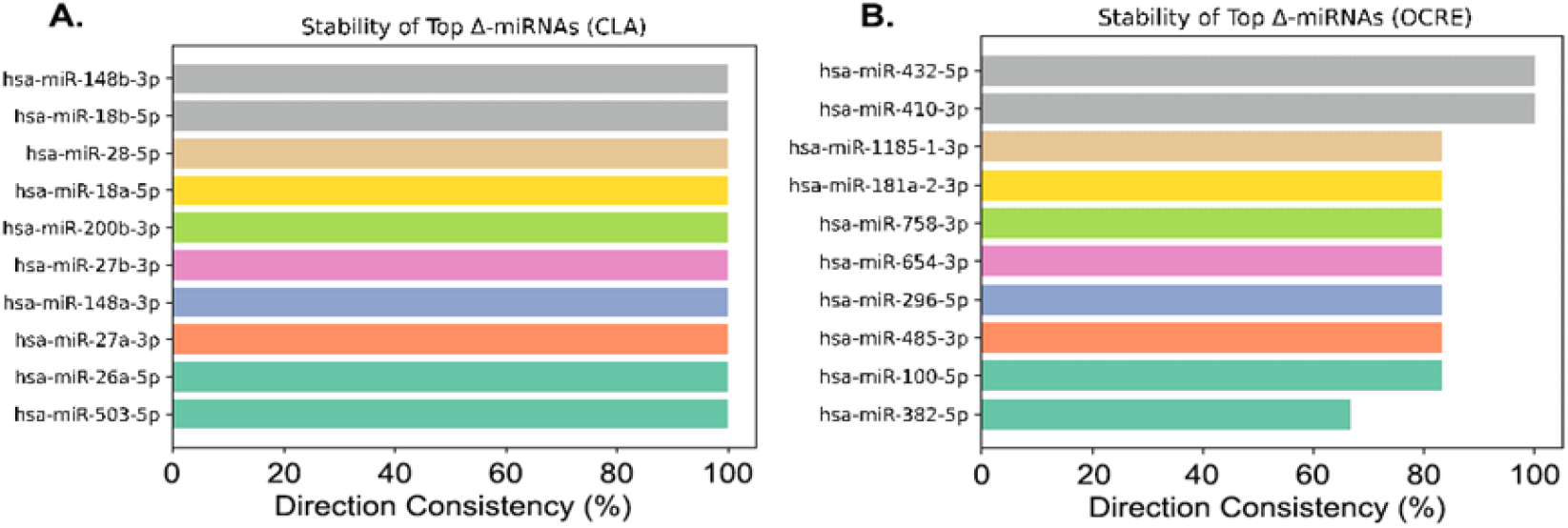
**(A)** Directional consistency of top-ranked Δ-miRNAs following cladribine treatment, expressed as the percentage of patients exhibiting Δ-expression in the same direction as the group mean. **(B)** Directional consistency of top-ranked Δ-miRNAs following ocrelizumab (OCRE) treatment. Bars indicate the proportion of subjects showing concordant directionality for each miRNA.

On the other hand, stability patterns of Δ-miRNAs linked to ocrelizumab therapy were more variable **(Figure 4B)**. More inter-individual heterogeneity was reflected by the moderate consistency of certain miRNAs across patients, whereas others exhibited high directional consistency. This trend is consistent with ocrelizumab’s more focused and specific molecular actions compared to other anti-CD20 therapies(Cree, Berger, & Greenberg, 2025). Compact and treatment-specific miRNA signatures for cladribine and ocrelizumab (OCRE) were obtained by integrating statistical evidence, ranking, and stability measures. These miRNAs are the most repeatable longitudinal signals found throughout the analytical process(Dong et al., 2013).

### Functional Consequences of miRNA changes

We noted unique miRNA’s not identified by (Arisi et al., 2023) following either ocrelizumab or cladribine therapy. Many are related to MS and a few are known to be elevated in progressive forms of the disease as depicted in Table 6.

**Table 6:** Overview of potential cellular targets: Summarizes potential miRNA cellular targets unique to either OCRE or CLA. Only hsa-miR-27b-3p was common between the prior analysis by Arisi et al and our current t0- and t1-paired analysis.

| Treatment | miRNA | Validated/Predicted Targets | Function in relation to MS | Citations |
| --- | --- | --- | --- | --- |
| CLA | hsa-miR-503-5p | CDC25a, CCND1, BCL2 | Not directly implicated in MS, Regulation of cell cycle and apoptosis under stress. | (Sarkar, Dey, & Dutta, 2010)<br>(M. Zhang et al., 2025) |
| CLA | hsa-miR-26a-5p | SLC1A1<br>Glutamate receptor signaling; inversely correlated to DLG4 | Upregulated in Natalizumab and IFNβ treatment | (Potenza et al., 2018)<br>(Mameli et al., 2016),<br>(Bruna De Felice et al., 2014) |
| CLA | hsa-miR-27a-3p | PPARγ, Wnt3a, Wnt-β-catenin signaling pathway modulator | Th17/Treg balance, OPC differentiation, Linked to RRMS=> SPMS transition | (Kulyté et al., 2019)<br>(Tripathi A et al., |
|  |  |  |  | 2019)<br>(Regev K et al., 2016) |
| CLA | hsa-miR-148a-3p | HLA-G, DNMT1 | Epigenetic regulation of immune tolerance, Low serum levels linked to primary progressive MS | (Manaster et al., 2012)<br>(Muñoz-San Martín M et al., 2023) |
| CLA | hsa-miR-27b-3p | PPARγ, FOXO1 | Impacts lipid metabolism and Th1/Th2 balance | (Kulyté et al., 2019)<br>(D'Onofrio N et al., 2023)<br>(Guerau-de-Arellano M et al., 2011) |
| OCRE | hsa-miR-100-5p | mTOR, IGF1R | Not directly implicated in MS, Modulates autophagy and cell survival. | (Liu H et al., 2024)<br>(Pinheiro AHG et al., 2024) |
| OCRE | hsa-miR-410-3p | STAT3, ELAVL4 | Reduced in RRMS, Impairs pro-inflammatory cytokine secretion | (Liu D et al., 2016)<br>(Keller A et al., 2015)<br>(Wang Y et al., 2019) |
| OCRE | hsa-miR-432-5p | ADAR1, Syt7 | Elevated in gray matter lesions and in progressive MS patients, Regulation of RNA editing and innate immune sensing. | (Fritsche L et al., 2020)<br>(Vistbakka J et al., 2017)<br>(Zhang X et al., 2021) |
| OCRE | hsa-miR-296-5p | IKBKE, NumbL, STAT5a | Regulated by circular RNA linked to MS, Modulates NF-κB signaling | (Li L et al., 2022)<br>(Li H et al., 2018) |
| OCRE | hsa-miR-485-3p | PGC-1α, STAT3 | Downregulated in RRMS compared to control patients, Regulates Th17 response in experimental autoimmune encephalitis, Mitochondrial biogenesis and energy homeostasis. | (Alizadeh-Fanalou et al., 2020)<br>(Ryu et al., 2026)<br>(Koh et al., 2021) |

## Discussion

Comprehensive quality control analyses confirmed that the miRNA expression dataset exhibits high technical integrity and is suitable for downstream longitudinal and treatment-specific analyses. The lack of extreme outliers in unsupervised principal component analysis and density **(Figure 1A-B)** and the strong overlap seen in sample-wise density distributions show that global expression patterns are very consistent across samples and are not dominated by batch-driven effects or technical artifacts. The observed variability appears to be mostly due to biological heterogeneity rather than noise, as this consistency is consistent with the Agilent microarray workflow’s efficient platform-level preprocessing and normalization(Kahraman et al., 2018). Importantly, the QC analyses’ absence of noticeable non-biological factor separation or clustering lends credence to the reliability of later within-subject comparisons and treatment-stratified analyses, although this behaviour is more indicative of technological resilience than biological signal loss. When taken as a whole, these QC results give assurance that the dataset provides a stable and trustworthy basis for examining long-term changes in miRNA expression linked to disease-modifying treatments, supporting the move to paired analyses that concentrate on treatment-induced molecular dynamics.

### Longitudinal Pairing and Sample Structure

The rigorous criterion for full t0–t1 pairing improves statistical robustness by allowing direct within-subject comparisons, therefore decreasing confounding from inter-individual variability, even when the number of matched patients is limited. Internal validation of measurement consistency across timepoints and further support for data dependability are provided by the existence of several technical replicates for various patients(Klebanov & Yakovlev, 2007). When taken as a whole, the paired sample structure shown in **Table 3** provides a solid basis for further studies that concentrate on changes in treatment-associated miRNA expression and supports the use of longitudinal difference-based methods.

The Δ-miRNA study demonstrated unique and treatment-specific molecular response patterns subsequent to cladribine and ocrelizumab therapy. **Figure 2A** demonstrates that cladribine therapy resulted in a coordinated alteration in miRNA expression, with several miRNAs showing significant positive Δ-expression and uniform directionality among patients. The inclusion of many miRNAs from related families among the most significant signals **(Table 4)** indicates organized regulatory alterations rather than random or isolated effects, aligning with the extensive immunomodulatory and immunological reconstitution processes associated with cladribine. Conversely, the Δ-miRNA profile after ocrelizumab therapy exhibited a more subdued pattern of alteration **(Figure 2B)**, with a reduced number of miRNAs demonstrating significant changes in expression. This selective response likely indicates the specific mechanism of B-cell depletion and the anticipation of more nuanced downstream transcriptional regulation, as previously shown to be the case for B cell differentiation (Laidlaw & Cyster, 2021). Although the quantity of change is diminished, the discernible directionality and consistency among the predominant miRNAs suggest that ocrelizumab medication is still linked to certain miRNA-level modifications.

Collectively, our data illustrate that cladribine and ocrelizumab elicit unique miRNA response signatures, underscoring therapy-specific molecular reconfiguration rather than a common therapeutic impact. The miRNAs listed in **Table 4** constitute a targeted selection of candidates for subsequent integrative studies and endorse the application of Δ-based, within-subject methodologies to elucidate treatment-related molecular dynamics. These results establish a molecular framework for subsequent stability-aware and machine-learning–based analyses aimed at characterizing treatment response patterns. The miRNAs found in this research most likely represent immune remodelling mechanisms linked to therapy that take place in response to multiple sclerosis disease-modifying therapies. A number of the cladribine-associated miRNAs have previously been connected to inflammatory signalling pathways and immunological control. For instance, miR-503-5p has been linked to the control of inflammatory signalling networks and immune cell proliferation, indicating that its elevated expression may represent immunological reconstitution activities after cladribine-induced lymphocyte depletion. Similar to this, miR-26a-5p has been shown to control cytokine signalling pathways and T-cell activation, indicating that it may play a part in influencing adaptive immunological responses during treatment-associated immune rebalancing. The coordinated overexpression of members of the miR-27 family, such as miR-27a-3p and miR-27b-3p, which are recognized regulators of inflammatory responses and macrophage polarization, may suggest more extensive modulation of innate immune signalling after cladribine treatment. The miRNA profile seen after ocrelizumab therapy, on the other hand, is more selective and contains miR-100-5p, a regulator that was previously linked to immune regulatory pathways and B-cell development. This finding is in line with ocrelizumab’s mode of action, which modifies downstream immune signalling networks by specifically eliminating CD20-positive B cells. When combined, these findings imply that the miRNA signatures found in this study may represent therapy-specific transcriptional regulatory programs that reflect different immune modulation mechanisms brought on by ocrelizumab-mediated B-cell depletion and cladribine-mediated immune reconstitution. These miRNAs are physiologically reasonable candidates for further research aiming at comprehending molecular responses to disease-modifying medications in multiple sclerosis, even though functional validation will be necessary.

The ranking analysis refined the Δ-miRNA signals identified in earlier analyses into treatment-specific molecular signatures. Data in **figure 3A** illustrates that cladribine therapy resulted in a significant and coordinated alteration in the expression of a specific group of miRNAs, along with extensive immunomodulatory and immune-reconstitution effects. The recurrence of miRNAs from related families among the highest-ranked candidates **(Table 4)** indicates the existence of organized regulatory responses rather than random fluctuation(Ebert & Sharp, 2012). Conversely, ocrelizumab therapy yielded a more subdued ranking profile **(Figure 3B)**, marked by a reduced number of miRNAs exhibiting moderate effect sizes. This pattern corresponds with the intended mode of action of ocrelizumab and indicates selective downstream transcriptional regulation. The found ranking miRNAs for ocrelizumab offer insight into certain biological pathways activated post-therapy(Wei et al., 2025). The graded Δ-miRNA profiles indicate that cladribine and ocrelizumab elicit unique and treatment-specific miRNA response patterns.

### Interpretation of stability findings

The stability analysis presented provides critical validation of the treatment-associated Δ-miRNA signals identified in earlier results. The idea that these variations represent a common molecular response rather than individual-specific fluctuations is supported by the high directional consistency of cladribine-associated miRNAs across patients, as illustrated in **(Figure 4A)**. The wide-ranging immunomodulatory and immunological reconstitution effects linked to cladribine treatment are in line with this stability(Alroughani et al., 2025; Lünemann, Ruck, Muraro, Bar-Or, & Wiendl, 2020) The ocrelizumab-associated miRNA profile, on the other hand, shows significant inter-individual variability **(Figure 4B)**, indicating a more varied molecular response. This result suggests that downstream transcriptional effects may differ across individuals and is consistent with the targeted mechanism of B-cell depletion. Crucially, despite their decreased overall stability, a number of miRNAs continue to show strong directional consistency, confirming their significance as treatment-associated signals. The final treatment-associated Δ-miRNA signatures identified in this study represent the most reproducible longitudinal molecular alterations observed across the analytical workflow. These signatures capture miRNAs that consistently respond to therapy instead of showing random variation or isolated statistical significance(Navon et al., 2009). They do this by combining longitudinal change, effect magnitude, and directional stability across individuals. The cladribine-associated signature is defined by precisely coordinated miRNA regulation exhibiting consistently high directional stability among patients, signifying a collective molecular response to treatment. This pattern aligns with the extensive immunomodulatory and immune-reconstitution effects of cladribine, reinforcing the notion that these miRNAs represent authentic treatment-related regulatory mechanisms. In contrast, the signature associated with ocrelizumab shows stable but more varied longitudinal modulation, which has also been noted in human T cells in response following Ocrelizumab treatment Although ocrelizumab-associated Δ-miRNAs exhibit greater inter-individual variability than cladribine-associated signals, several miRNAs still demonstrate consistent directional changes across patients. This suggests that the therapy induces selective but reproducible transcriptional modulation, rather than the broader coordinated response observed following cladribine treatment. These results show that strong changes in miRNA levels that are linked to treatment can be found without relying only on isolated statistical significance. By focusing on longitudinal consistency and reproducibility we developed a biologically-based way to understand therapy-specific molecular responses and a clear list of candidate miRNAs for future validation studies.

## Conclusion

This study conducted a longitudinal examination of treatment-related miRNA dynamics in MS by within-subject Δ-miRNA profiling. Through the integration of statistical testing, effect-size ranking, and stability evaluation across people, we discerned consistent and treatment-specific miRNA response patterns subsequent to cladribine and ocrelizumab medication. The findings indicate that cladribine is linked to a highly coordinated miRNA response with significant directional consistency among patients, while ocrelizumab elicits a more selective and variable miRNA modulation pattern. The definitive miRNA signatures presented in **Table 4** underscore molecular alterations that are consistent among people and unlikely to be influenced by individual-specific influences. This work prioritizes repeatability and longitudinal consistency above isolated statistical significance as essential criterion for identifying physiologically relevant treatment-associated miRNA signals. These findings collectively offer a consistent molecular framework for comprehending therapy-specific immune modulation in MS and build a solid platform for future translational and mechanistic research.

### Future Scope

Multiple avenues can enhance and reinforce the conclusions of this research. Initially, the validation of the discovered miRNA signatures in larger, independent cohorts is crucial to ascertain their generalizability across varied patient groups and therapeutic contexts. The inclusion of extended follow-up periods may elucidate the temporal stability of these miRNA responses and their correlation with prolonged therapy benefits. Secondly, the integration of the discovered miRNAs with downstream mRNA, proteomic, or immune-cell-specific data may clarify the molecular pathways by which these miRNAs influence treatment response. Multi-omics methodologies might enhance the biological understanding of therapy-specific regulation networks. Subsequently, future research may investigate the therapeutic significance of these miRNA signatures by analysing correlations with therapy efficacy, illness progression, or long-term results. With proper validation, persistent longitudinal miRNA signatures may function as relevant biological markers to assist in treatment monitoring and tailored therapeutic approaches in MS. This study has several limitations. First, the number of patients with complete longitudinal pairs was small (n=4 for cladribine and n=6 for ocrelizumab), which limits statistical power. Second, the analysis relied on publicly available microarray data without access to detailed clinical outcomes, preventing correlation of miRNA changes with treatment response or disease activity. Third, only a single post-treatment timepoint (6 months) was available, limiting evaluation of temporal dynamics. Finally, functional validation of the identified miRNA candidates was beyond the scope of this study.

### Source of Data

The miRNA expression data analysed in this study were obtained from the NCBI GEO # accession number GSE230064, a publicly accessible repository hosting high-throughput functional genomics datasets. The dataset comprised longitudinal miRNA expression profiles from MS patients treated with cladribine or ocrelizumab, including paired baseline and post-treatment samples. The data were downloaded from GEO, curated, and reanalysed locally for the purposes of this study.

